# *Ex situ* spawning, larval development, and settlement in the massive reef-building coral *Porites lobata* in Palau

**DOI:** 10.1101/2024.08.19.608627

**Authors:** Matthew-James Bennett, Carsten G.B. Grupstra, Jeric Da-Anoy, Maikani Andres, Daniel Holstein, Ashley Rossin, Sarah W. Davies, Kirstin S. Meyer-Kaiser

**Author notes:** Correspondence: Woods Hole Oceanographic Institution, 266 Woods Hole Road, Woods Hole, MA 02543 USA.

## Abstract

Reproduction, embryological development, and settlement of corals are critical for survival of coral reefs through larval propagation. Yet, for many species of corals, a basic understanding of the early life-history stages is lacking. In this study, we report our observations for *ex situ* reproduction in the massive reef-building coral *Porites lobata* across two years. Spawning occurred in April and May, on the first day after the full moon with at least two hours of darkness between sunset and moonrise, on a rising tide. Only a small proportion of corals observed had mature gametes or spawned (17 – 35%). Eggs were 185 – 311 μm in diameter, spherical, homogenous, and provisioned with 95 – 155 Symbiodiniaceae algae. Males spawned before females, and *ex situ* fertilization rates were high for the first 2 hours after egg release. *P. lobata* larvae were elliptical, approximately 300 μm long, and symbiotic. Just two days after fertilization, many larvae swam near the bottom of culture dishes and were competent to settle. Settlers began calcification two days after metamorphosis, and tentacles were developed 10 days after attachment. Our observations contrast with previous studies by suggesting an abbreviated pelagic larval period in *P. lobata*, which could lead to the isolation of some populations. The high thermal tolerance and a broad geographic range of *P. lobata* suggest this species could locally adapt to a wide range of environmental conditions, especially if larvae are locally retained. The results of this study can inform future work on reproduction, larval biology, dispersal, and recruitment of *P. lobata*, which could have an ecological advantage over less resilient coral species under future climate change.

## 1. Introduction

Sexual reproduction, larval dispersal, and recruitment are key processes that are essential for the long-term maintenance of coral reefs in the face of myriad threats (Hughes & Tanner 2000, Hughes et al. 2017, Richmond et al. 2018). Sexually-produced larvae can replenish coral populations with genetically-variable individuals to withstand present selective pressures.

Environmental stressors such as elevated temperatures can result in reduced reproductive capacity in corals and a breakdown in the influx of new coral colonies to replace those that suffer mortality, driving down the abundance and genetic diversity of corals on a reef (Hagedorn et al. 2016; Fisch et al. 2019, Henley et al. 2022, Humphrey et al. 2008). Improving our understanding of the fundamental life history stages and key functions of reef-building corals that drive their population dynamics, such as their sexual reproduction, is essential as efforts to manage coral reefs globally expand (Edwards et al. 2024).

Maintaining species and genetic diversity in coral populations is vital to ensuring coral reef longevity and the long-term success of restoration initiatives (Mcleod et al. 2019, Shaver et al. 2022). Recent and past studies on coral reproduction have concentrated on a limited range of species, growth forms and reproductive modes, with 30% of recent studies involving hermaphroditic branching corals of the *Acropora* genus (Boström-Einarsson et al. 2020). This limited focus presents a challenge for predicting changes in a reef’s community structure.

Recent work has emphasized the importance of incorporating corals with diverse life histories into reproductive studies and restoration strategies. However, basic information on reproductive biology is missing for many species, making their inclusion in such studies more challenging (Guest et al. 2023). The ability to predict the timing of reproductive events *in situ*—which can vary across the geographic range of a given species— requires extensive observational data to allow practitioners and managers to effectively plan research operations and limit activities that may disrupt this important natural process (Kenyon 1995, Marhaver et al. 2015, Baird et al., 2009, Baird et al. 2022).

Active coral reef restoration approaches that utilize sexual propagation through assisted fertilization and recruitment have the potential to repopulate degraded reefs (dela Cruz & Harrison 2017). These interventions are increasingly stressed as an important strategy to promote the survival of coral reefs globally in addition to reducing greenhouse gas emissions and informed ecosystem management plans (Kleypas et al. 2021, Vardi et al. 2021, Suggett et al. 2024). However, the efficacy and productive yield of these initiatives is limited by the lack of available information for many coral species (Boström-Einarsson et al., 2020; Guest et al., 2023).

In corals, sexual reproduction presents a diverse range of strategies and reproductive modes, even within broadcast spawning species (Guest et al. 2012). Most *ex situ* spawning studies or restoration efforts have concentrated on hermaphroditic species because the gamete bundles produced by these species, which encase both eggs and sperm, facilitate the identification of fecund adult colonies and the collection and concentration of gametes (Boström-Einarsson et al. 2020). In gonochoric species where sperm is released directly into the water column, collection of highly concentrated sperm can be more challenging as the sperm dilutes on release. Lower sperm concentrations can result in lower *in vitro* fertilization success (Nozawa et al. 2015, dela Cruz & Harrison 2020). Sperm limitation has also been proposed as a factor leading to reduced natural fertilization rates as reefs degrade and adult colonies become rarer and more dispersed (Levitan & Petersen 1995). In response, cryopreservation of sperm has been used in *in vitro* fertilization efforts to boost genetic diversity of corals and promote desirable genotypes (e.g., Grosso-Becerra et al. 2021). External fertilization success in corals can additionally be impacted by the specific timing after release when compatible gametes encounter each other, and the duration that they remain in contact (dela Cruz & Harrison 2020).

*Porites lobata* is a gonochoric broadcast spawning coral (i.e., with separate male and female colonies) that is widely distributed and abundant in the Pacific Ocean (Glynn et al. 1994, Baums et al. 2012, Sale et al. 2019). It is a dominant reef builder, forming massive mounding colonies, and has demonstrated relatively high tolerance to anthropogenic stressors (Loya et al. 2001, Levas et al. 2013, Barshis et al. 2018). A recent study identified distinct genetic lineages within the *Porites* cf. *lobata* species complex across habitat gradients in the low-latitude reefs of Palau, with some lineages exhibiting elevated thermal tolerance (Rivera et al. 2022). However, little is known about the early life-history stages in this population of *P. lobata*. Previous studies show differences in the timing of spawning events across the species’ geographic range, leading to questions about the relevant environmental cues (Table 1). Only one study has reported direct, *in situ* observations of gamete release in Palau, with a second based on histological observations (Penland et al. 2004, Gouezo et al. 2020).

**Table 1:** Reproductive timing of *Porites lobata* reported in various geographic locations. dAFM, days after full moon; PICR, Palau International Coral Reef Center. Note that many observations are from histological studies where the night of spawning could not be accurately determined. All other studies are spawning observations denoted as *in situ* or *ex situ*.

| Country | Site | Year | Month | dAFM | Observation type | References |
| --- | --- | --- | --- | --- | --- | --- |
| Australia | Heron Island | 1978-79 | Dec | N/A | Histology | Kojis & Quinn, 1981 |
| Australia | Orpheus Island | 1983 | Nov | 5 | *In situ* | Babcock et al., 1986 |
| Chile | Rapa Nui | 2013-14 | Feb, Mar | N/A | Histology | Buck-Wiese et al., 2018 |
| Costa Rica | Caño Island | 1985-91 | Apr, May, Sep, Oct | N/A | Histology | Glynn et al., 1994 |
| Ecuador | Santa Cruz Island | 1985-91 | Jan, Apr, May, Sep, Oct | N/A | Histology | Glynn et al., 1994 |
| Indonesia | Karimunjawa | 1995 | Oct | 3 | *In situ* | Tomascik et al., 1997 |
| Japan | Sesoko Islands | 1986 | Jun | 2 | *In situ* | Isomura & Fukami, 2018 |
| Palau | Nikko Bay | 2002-03 | Apr, May | 3-4 | *In situ* | Penland et al. 2004 |
| Palau | PICRC | 2017-18 | Apr, May | N/A | Histology | Gouezo et al. 2020 |
| Palau | Chelbacheb | 2022 | Apr, May | 3-4 | *Ex situ* | This study |
| Palau | Chelbacheb | 2023 | Apr, May | 2-4 | *Ex situ* | This study |
| Panama | Uva Island | 1985-91 | Feb, Sep | N/A | Histology | Glynn et al., 1994 |
| Panama | Taboga Island | 1985-91 | Jul, Aug | N/A | Histology | Glynn et al., 1994 |
| Saudi Arabia | Al Fahal | 2012 | May | 1 | *In situ* | Bouwmeester et al., 2015 |
| USA (Hawai‘i) | Kāne‘ohe Bay | 1987 | Aug, Sep | 3-6 | Histology | Hodgson, 1988 |
| USA (Hawai‘i) | Kāne‘ohe Bay | 1997 | Jun, Jul, Aug | 1-4 | *Ex situ* | Field, 1998; Maté, 1998 |

Given the broad geographic distribution of *P. lobata* across diverse habitat types, demonstrating a high degree of plasticity and potential for local adaptation, the evidence of relatively high resilience to thermal stress, and the ecological importance of this species in coral reef systems, *P. lobata* represents a promising candidate for restoration initiatives (Humanes et al. 2021). In this study, we expand the body of knowledge on reproduction and embryological development in *P. lobata*. We report histological observations of gametogenesis, direct observations of *ex situ* spawning, describe the release of eggs and sperm, assess fertilization with gametes at varying times after release to determine the window for optimal fertilization success, and track larval development as well as settlement behaviour, metamorphosis, and development in the first few weeks after settlement. We conducted initial trials with established methods for cryopreservation of coral sperm, which yielded limited success in this species (see Supplementary Material). With this study, we aim to provide detailed descriptions and a baseline record of the reproductive biology and early life stages of *P. lobata,* which can enhance the capacity for further research on this species, and aid its integration into restoration programmes employing sexual coral propagation as a tool for coral reef conservation.

## 2. Methods

### 2.1 Environmental data and colony collection

*Porites lobata* colonies (minimum diameter 15 cm) were collected from our six study sites (n = 7–30 per site) in the Rock Islands Southern Lagoon (Chelbacheb) in the Republic of Palau (Figure 1, Supplementary Material). Colonies were collected around the full moon, from 2 days before to 4 days after the full moon (dAFM). For smaller colonies, (15–25 cm diameter), the whole colony was collected; for larger colonies (>25 cm diameter), a piece with diameter 15–25 cm was broken off using a hammer and chisel. Corals were housed in individual plastic containers (4.2 L; 17 x 13 x 19 cm) and distributed across flow-through aquariums (3000 L) at the Palau International Coral Reef Center (PICRC) during the observation period. Colonies were returned to nurseries at their site of collection between observation periods so that the same colonies could be repeatedly observed. In 2023, additional colonies were collected from Risong, Taoch and Outer Taoch sites, from which spawning had been observed in 2022, to increase the likelihood of observing gamete release (Supplementary Material).

**Figure 1.**
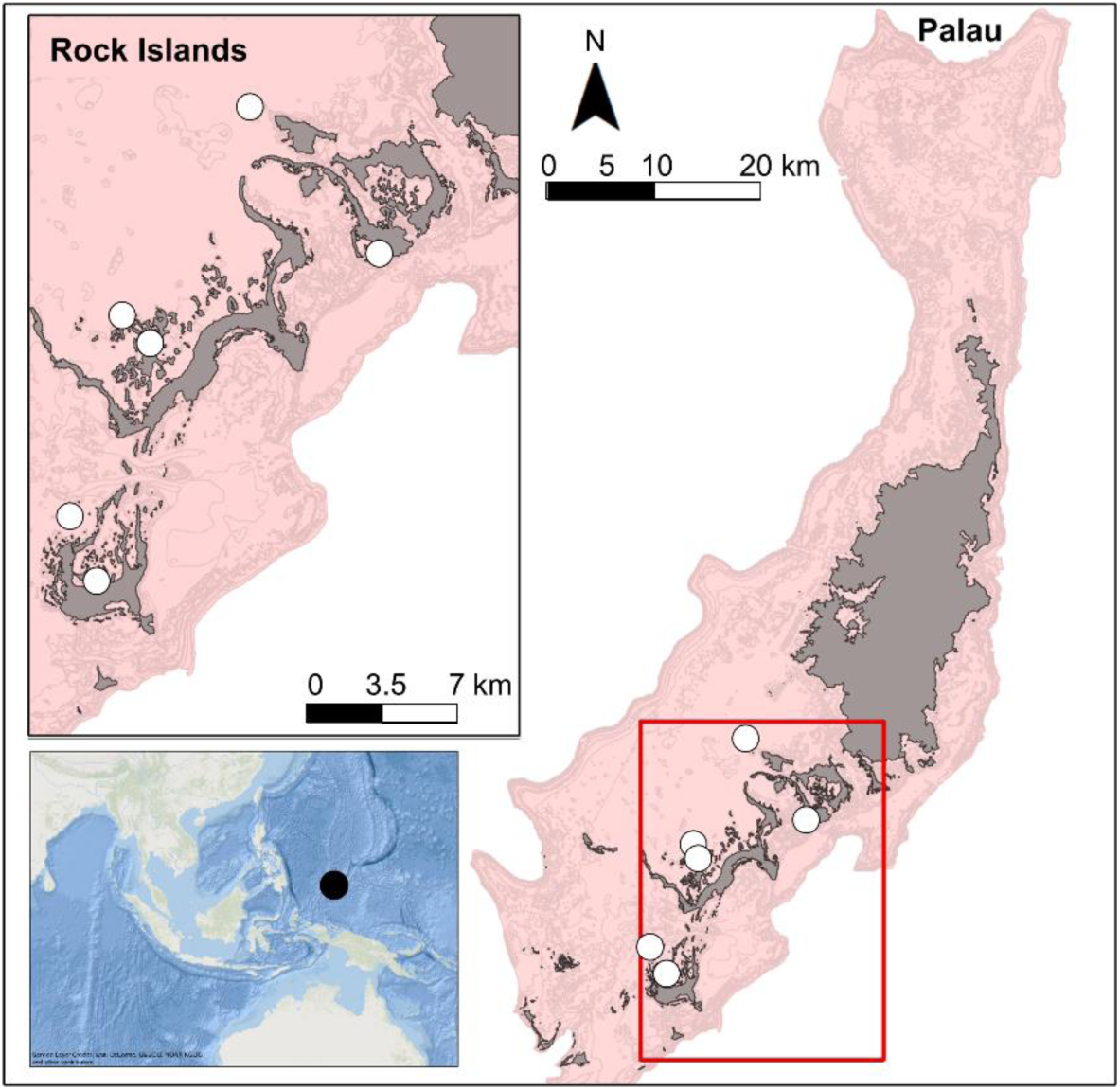
Coral collection sites in the Rock Island Southern Lagoon (Chelbacheb), Palau. Pink indicates the reef platform, while grey represents land.

Environmental factors that may serve as cues for synchronised gamete release were recorded *in situ* or sourced from publicly available datasets. Water temperatures were recorded using loggers at ∼3 m depth (HOBO TidbiT v2, Onset Corp.) at our six study sites November 2021 – May 2023. Temperature measurements were averaged across all sites for each lunar month. The first lunar month of the year begins on the first full moon following the winter solstice, and each lunar month constitutes one lunar cycle (Baird et al. 2022). Tidal amplitude data were obtained from the University of HawaiʻiSea Level Center for the station at Malakal Island, Palau (7° 19.80’ N, 134° 27.28’ E; https://uhslc.soest.hawaii.edu/stations/?stn=007#levels). Sunset, moonrise, and twilight times for Koror, Palau were obtained from the U.S. Naval Observatory (https://aa.usno.navy.mil/data/RS_OneYear).

### 2.2 Histological observations

Small tissue samples (∼2 cm diameter) were collected from the same colonies included in the spawning observations with a hammer and chisel via SCUBA (November 2021) or in the laboratory at the PICRC after coral collection but prior to spawning (May 2022). Samples were collected from the center of each colony to avoid colony edges where gametes may not be present. Immediately following collection in the lab or the end of the SCUBA dive (i.e., within 20 minutes of collection), samples were fixed in a solution of 10% neutral buffered formalin in seawater.

Preserved samples were decalcified in 1% EDTA decalcifier solution (5% hydrochloric acid with 5.0 g EDTA L^-1^) for 24–48 hours and were stored in 70% ethanol after complete decalcification (Szmant-Froehlich et al. 1985, Glynn et al. 1991). Samples from November 2021 were processed and sectioned at the Seascape Ecology Lab at Louisiana State University, while the remaining samples were processed at to the Collaborative Research Laboratory (CoRe) at Boston University. Tissues were dehydrated, cleared, and paraffinized in a Leica ASP6025 Tissue Processor, embedded in wax blocks using a Leica EG1150H embedding machine, and then cooled in a freezer for 24 h before sectioning. Blocks were sectioned at 5 μm thickness in an oral-to-aboral direction at 300 μm intervals on a Leica RM2125RTS microtome. Three tissue sections for each sample were transferred on a microscope slide for staining.

Histological tissue sections were stained with hematoxylin and eosin or modified Heidenhain’s aniline on a Leica ST5020 multistainer. Sections from 43 individuals were examined for the presence or absence of male and female gametes under a binocular compound microscope (Olympus BH2) with a digital eyepiece camera attachment (Amscope). Gametes (oocytes and spermatocytes) were staged from I-V following the classification of Szmant-Froelich et al. (1985).

### 2.3 Spawning timing and gamete release

*Ex situ P. lobata* colonies were monitored in April and May 2022, November 2022, and April and May 2023 to observe spawning phenology and behaviour. On each observation day, the water level in aquariums was lowered to ∼15 cm shortly after sunset. Therefore, each coral was isolated in its own container during the observations, but containers were surrounded by running seawater so that the water remained at ambient ocean temperature. Spawning observations began < 1 hour after sunset and continued for 4 – 6 hours each day. All colonies were checked for gamete release every 10 – 20 minutes, and the frequency and time of gamete release was noted, as well as colony sex. Observations continued for 4 – 10 days in a given month.

### 2.4 Gamete collection and fertilization

Sperm were collected directly from colonies using a large plastic pipette during release, thereby maintaining high sperm concentrations, and transferred to 2.1 L hydrophobic plastic containers. Eggs were allowed to float to the surface so they could more easily be collected by pipette or by skimming the surface with a plastic tri-pour beaker or petri dish, before being transferred to 400 mL plastic containers with 5 µm filtered seawater (FSW). Gametes were kept separate until a sufficient number of parent colonies had spawned to begin a batch cross (minimum 3 males and 1 female). Sperm from selected male colonies were pooled in a single container, and eggs (from n=1 female per batch cross) were gently poured into this pool, allowing fertilization to commence. Gamete transfer was achieved by gently pouring only the eggs from the surface with as little seawater as possible to limit sperm dilution. For all crosses, gametes were separated 45 – 60 minutes after fertilization began. The surface layer with fertilized eggs was poured into clean plastic containers, eggs were rinsed with 5 µm FSW, and then eggs were gently split into multiple containers filled with 5 µm FSW to dilute any remaining sperm and lower the egg density.

In 2023, a subset of eggs was held apart from sperm until a set time after spawning in order to assess the impact of egg age on fertilization success. A small aliquot (∼10 mL) of concentrated sperm and ∼1000 eggs were added to triplicate wells of a 6-well plate for each time-point. Cell strainers (70 μm mesh) were used to separate eggs from sperm ∼45 minutes after fertilization began, and eggs were gently rinsed into a 50 mL beaker filled with FSW. Fertilization success was evaluated visually 2–3 hours after fertilization began by observing a sub-sample of the eggs from each cross under a dissecting microscope (Leica S9i) and counting whole (undivided) eggs and dividing embryos. In addition, we observed motility of a sub-sample of sperm periodically using a phase-contrast microscope (Amscope).

### 2.5 Embryological observations

Developing embryos and larvae were observed at least daily using a dissecting microscope and photographed using white light and blue light with a yellow filter (Nightsea) to detect fluorescence. Embryos and larvae were cultured in hydrophobic plastic containers (2.1 L) in still 5 µm FSW at ambient temperature (∼30° C), and water was changed every 1–2 days.

Egg size was estimated by measuring the diameter of 160 individual eggs using the straight-line selection tool in ImageJ. Densities of Symbiodinaceae algae were estimated in eggs and larvae using image analysis. Eggs and larvae were photographed in petri dishes under a dissecting microscope, and algal cells were counted in each individual using the cell counter function in ImageJ. Only a projection of the 3-dimensional surface area of each egg or larva was visible in the images, so the number of observed algae was used to extrapolate abundance per individual. The plan area of the egg or larva where algae could be clearly discerned in the image was measured using the oval select function. The diameter of the individual was measured using the straight-line function and used to calculate the 3-dimensional surface area of the egg or larva as 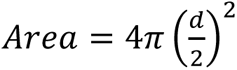. For elliptical larvae, two diameters were measured (long and short axes) and used to calculate surface area as 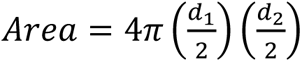. The number of algae observed was divided by the plan area and multiplied by the 3-dimensional surface area to yield the approximate number of cells in a larva. This image analysis method was compared to the standard method of quantifying algae from a known number of homogenized larvae using a haemocytometer. Three replicate solutions with 50 larvae homogenized in 100 µL FSW were prepared before adding 10 µL to the counting chamber. Both methods yielded similar ranges of algae per larva, so we report image analysis results for algae per egg and per larva in this study.

### 2.6 Settlement observations

Two days post fertilization, a sub-set of larvae were exposed to limestone tiles that had been pre-conditioned on the PICRC house reef and microscope slides with and without the addition of crushed crustose coralline algae (CCA, multiple species combined following Pollock et al. (2017)). Tiles were deployed horizontally on the seafloor on a shallow coral reef near PICRC for approximately one month for conditioning prior to settlement trials, but microscope slides were clean. Approximately 30 larvae were added to a 2.1 L hydrophobic plastic container with settlement substrata (tiles or glass slides). Settlers were observed daily and photographed using a dissecting microscope with white light and fluorescence.

## 3. Results

### 3.1 Histological observations

Histological analysis of *P. lobata* tissues revealed the presence of oocytes and spermatocysts containing spermatocytes in May 2022 but not in November 2021 (Figures 2A, 3). Out of the 43 colonies sampled in May, after the April and May 2022 spawning events, only 4.65% (2/43) colonies had oocytes and 13.95% (6/43) had spermatocysts (Figure 2A). The majority of the oocytes and spermatocysts were stage IV (Figure 3). The average size of mature (stage IV) oocytes was 186.98 ± 31.35 μm (mean ± stdev), and stage I-III oocytes were 83.12 ± 23.48 μm (Figure 2B). No hermaphroditic colonies (with both oocytes and spermatocysts) were found (Figure 3).

**Figure 2.**
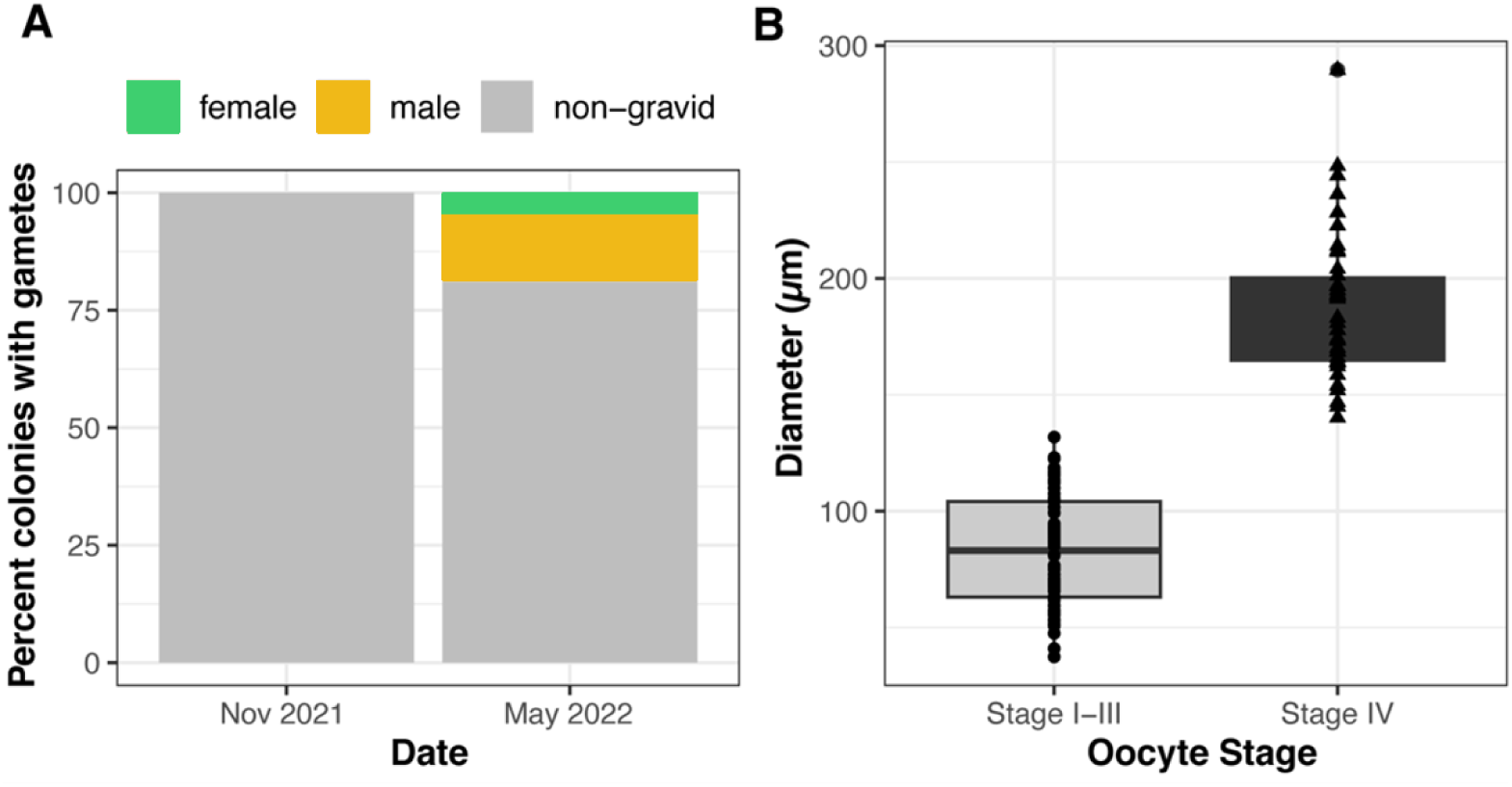
Percentage of gravid *Porites lobata* colonies and oocyte sizes. (A) Percentage of colonies containing oocytes (female) and spermatocysts (male) in each sampling date, n=43 individuals. (B) Immature (Stage I−III) and mature (Stage IV) oocyte sizes. Each point represents an oocyte measured in 10 random polyps of 2 gravid colonies found in May 2022.

**Figure 3.**
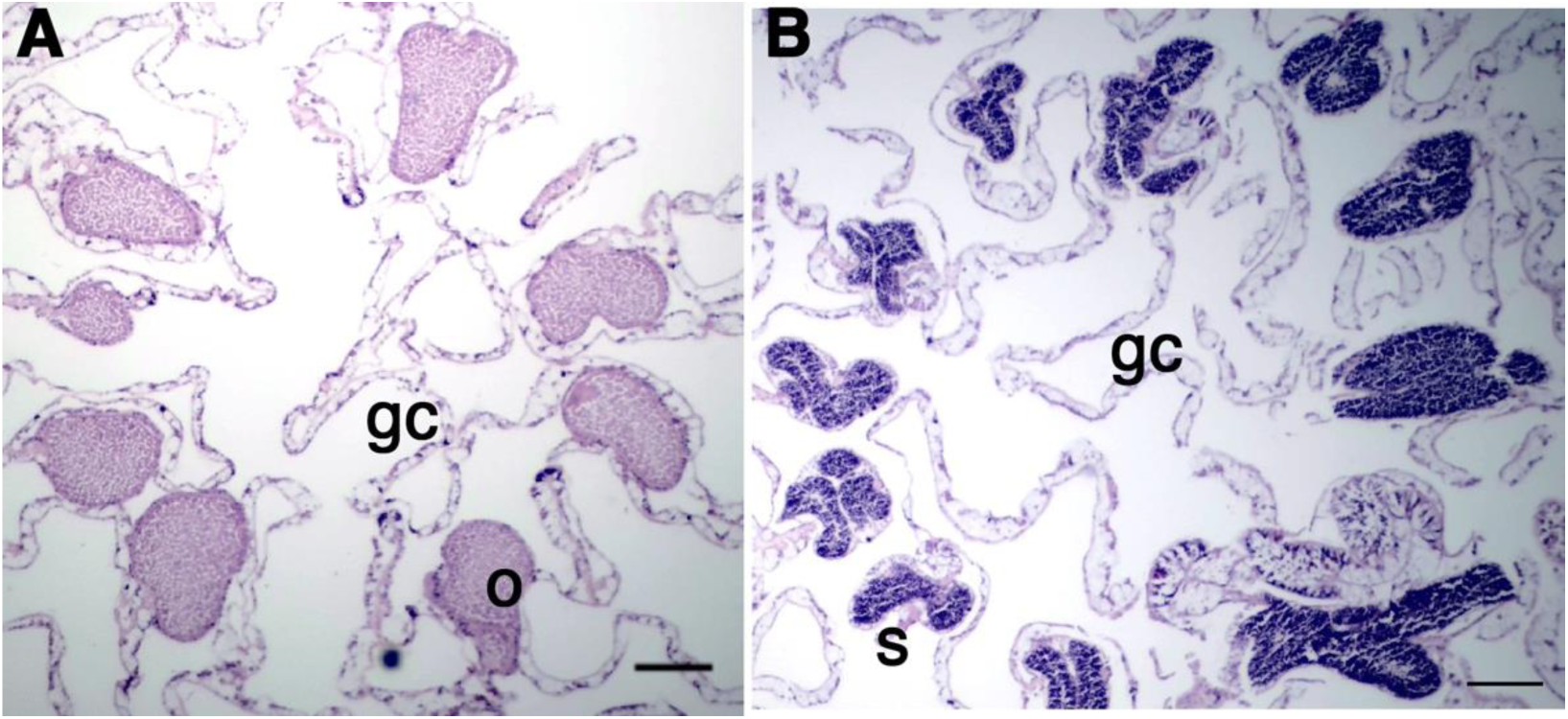
Photomicrographs of cross-sectional views showing gametes in *Porites lobata* tissues. (A) Histological section of a whole female polyp with oocytes. (B) Stage ІV spermatocysts. O, oocyte; S, spermatocyst; gc, gastrovascular cavity. Scale bars are 100 µm.

### 3.2 Spawning timing and gamete release

Spawning was observed 3–4 dAFM in April and May (Table 2). Gamete release times ranged 1–4 hours after sunset with the peak release (∼70% of spawning colonies) between 1.5 and 3.5 hours after sunset. Males began spawning earlier than females, and gamete release continued in pulses for both sexes for around 30 minutes. A small proportion of colonies (∼15%, 8/56) released gametes on more than one night in a given month.

**Table 2.** *Ex situ* spawning observations of *Porites lobata* colonies in Palau. Numbers in parentheses represent colonies that also spawned on the previous night. dAFM, days after full moon; mAS, minutes after sunset; No. colonies, total number of colonies observed in aquariums. Consecutive days with no spawning observed are condensed to a single line.

| dAFM | Date | No. colonies | No. spawning (repeat) males | Start of sperm release (mAS) | No. spawning (repeat) females | Start of egg release (mAS) |
| --- | --- | --- | --- | --- | --- | --- |
| 3 | Apr 20, 2022 | 34 | 4 | 90 – 258 | 2 | 190 – 320 |
| 4 | Apr 21, 2022 | 54 | 1 (1) | 58 | 3 (2) | 168 – 217 |
| 5-8 | Apr 22-24, 2022 | 54 | 0 | N/A | 0 | N/A |
| 1-3 | May 17-19, 2022 | 54 | 0 | N/A | 0 | N/A |
| 4 | May 20, 2022 | 54 | 1 | 198 | 1 | 172-210 |
| 5-9 | May 21-25, 2022 | 54 | 0 | N/A | 0 | N/A |
| 1-5 | Nov 10-14, 2022 | 54 | 0 | N/A | 0 | N/A |
| -2 to +2 | April 4-8, 2023 | 54 | 0 | N/A | 0 | N/A |
| 3 | April 9, 2023 | 89 | 4 | 149 – 199 | 3 | 179 – 229 |
| 4 | April 10, 2023 | 89 | 6 | 70 – 220 | 5 (1) | 170 – 335 |
| 5-8 | April 11-14, 2023 | 89 | 0 | N/A | 0 | N/A |
| -2 to +1 | May 4-7, 2023 | 98 | 0 | N/A | 0 | N/A |
| 2 | May 8, 2023 | 98 | 1 | 183 | 0 | N/A |
| 3 | May 9, 2023 | 98 | 6 | 139 – 209 | 6 | 159 – 276 |
| 4 | May 10, 2023 | 98 | 6 (3) | 69 – 149 | 8 (1) | 159 – 534 |
| 5-7 | May 11-13, 2023 | 98 | 0 | N/A | 0 | N/A |

In both 2022 and 2023, *P. lobata* spawning coincided with increasing water temperatures. Lunar months 3–5 each had higher mean water temperatures than the previous lunar month, and spawning occurred in lunar months 4 and 5 in both years (Figure 4). *P. lobata* spawning occurred exclusively on nights when moonrise was at least 2 hours after sunset, providing a significant dark period (Figure 5). Additionally, *P. lobata* spawned on a rising tide on nights when low tide occurred after sunset and high tide occurred on or after midnight (Figure 6).

**Figure 4.**
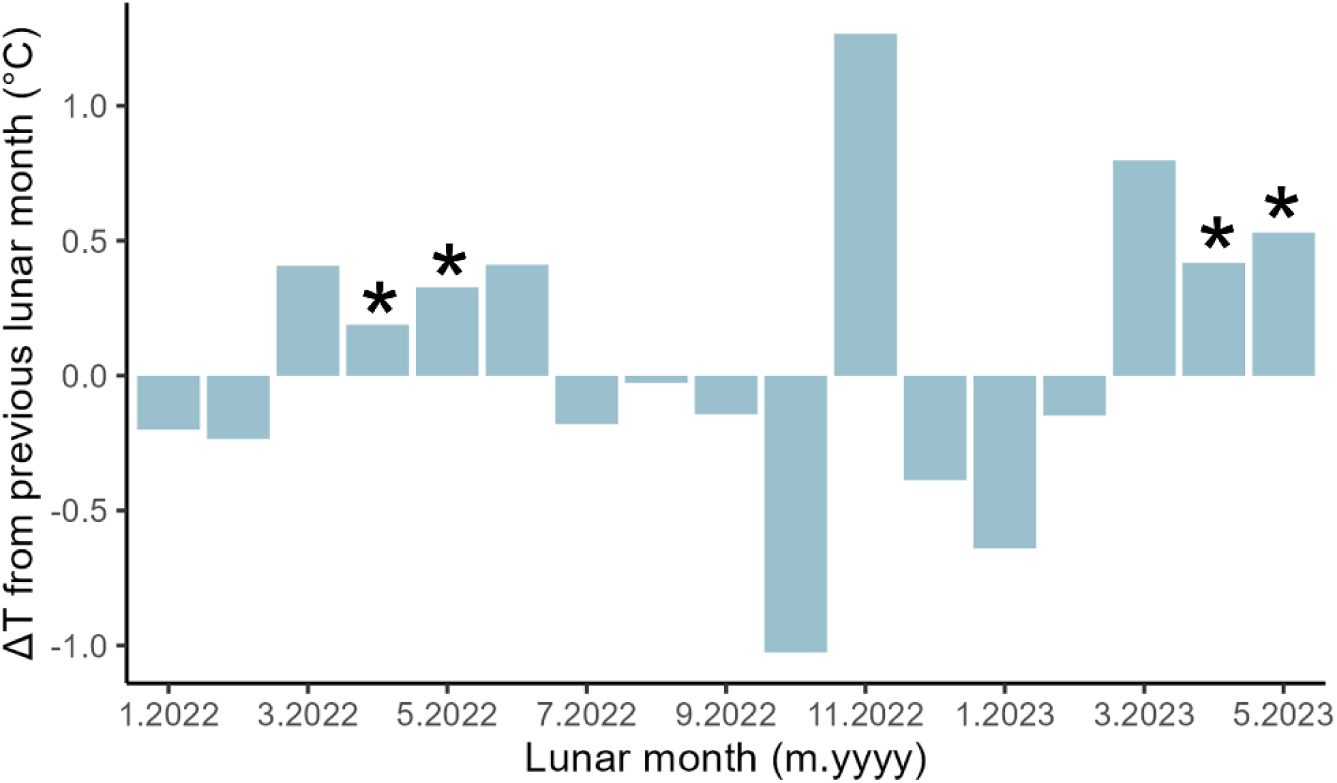
Monthly change in water temperature (°C) in *Porites lobata* habitats in Palau by lunar month. The first lunar month of the year begins with the first full moon following the winter solstice (Baird et al. 2022). Vertical bars show the difference in mean water temperature for a given lunar month compared to the previous lunar month. Asterix indicate lunar months in which *P. lobata* spawning was recorded. Observations were made in April and May 2022, November 2022 (no spawning observed), and April and May 2023.

**Figure 5.**
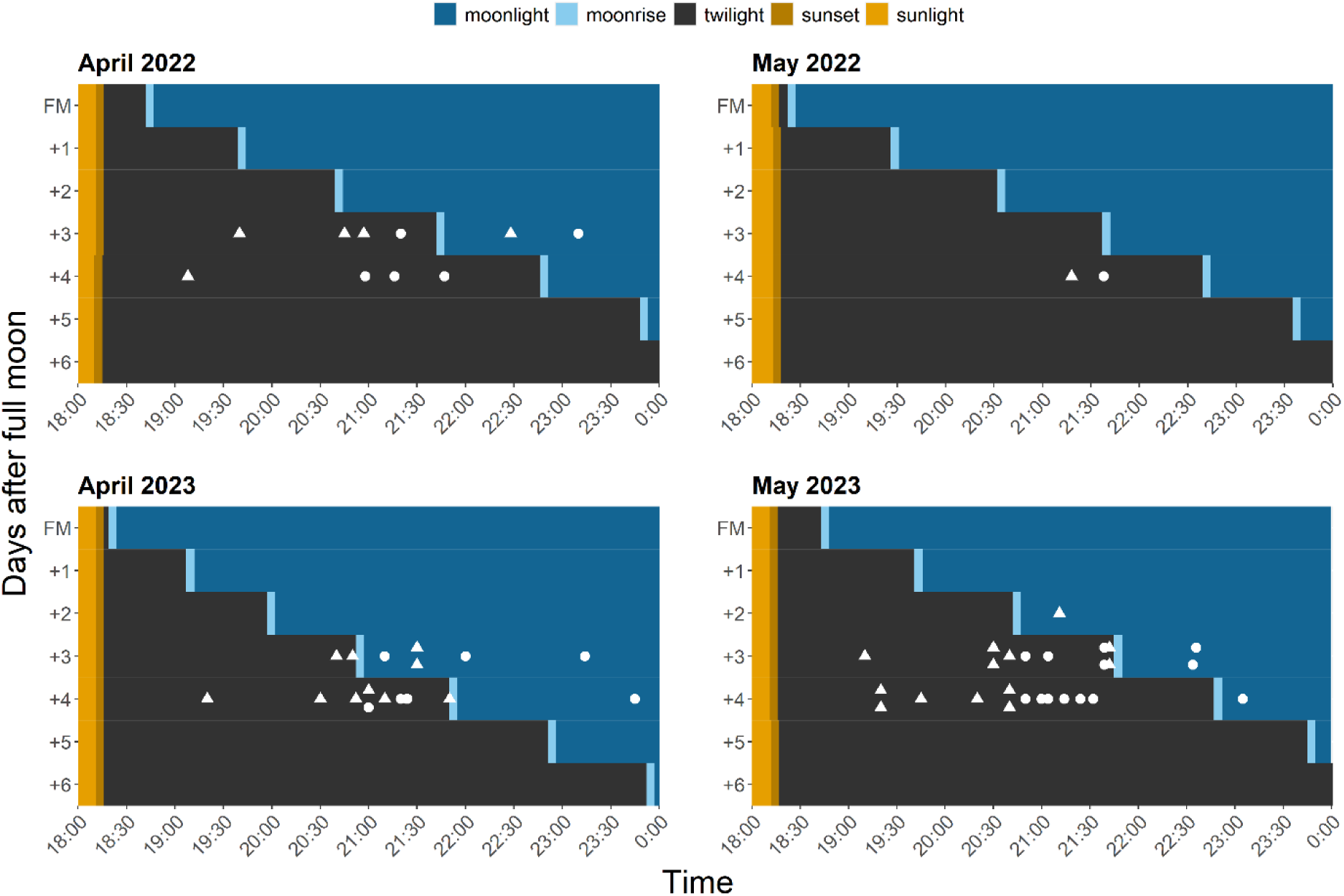
Spawning times for *Porites lobata* in relation to moonlight conditions in Palau. White shapes indicate observed gamete release times in each year of observation, with triangles representing male colonies and round points female colonies.

**Figure 6.**
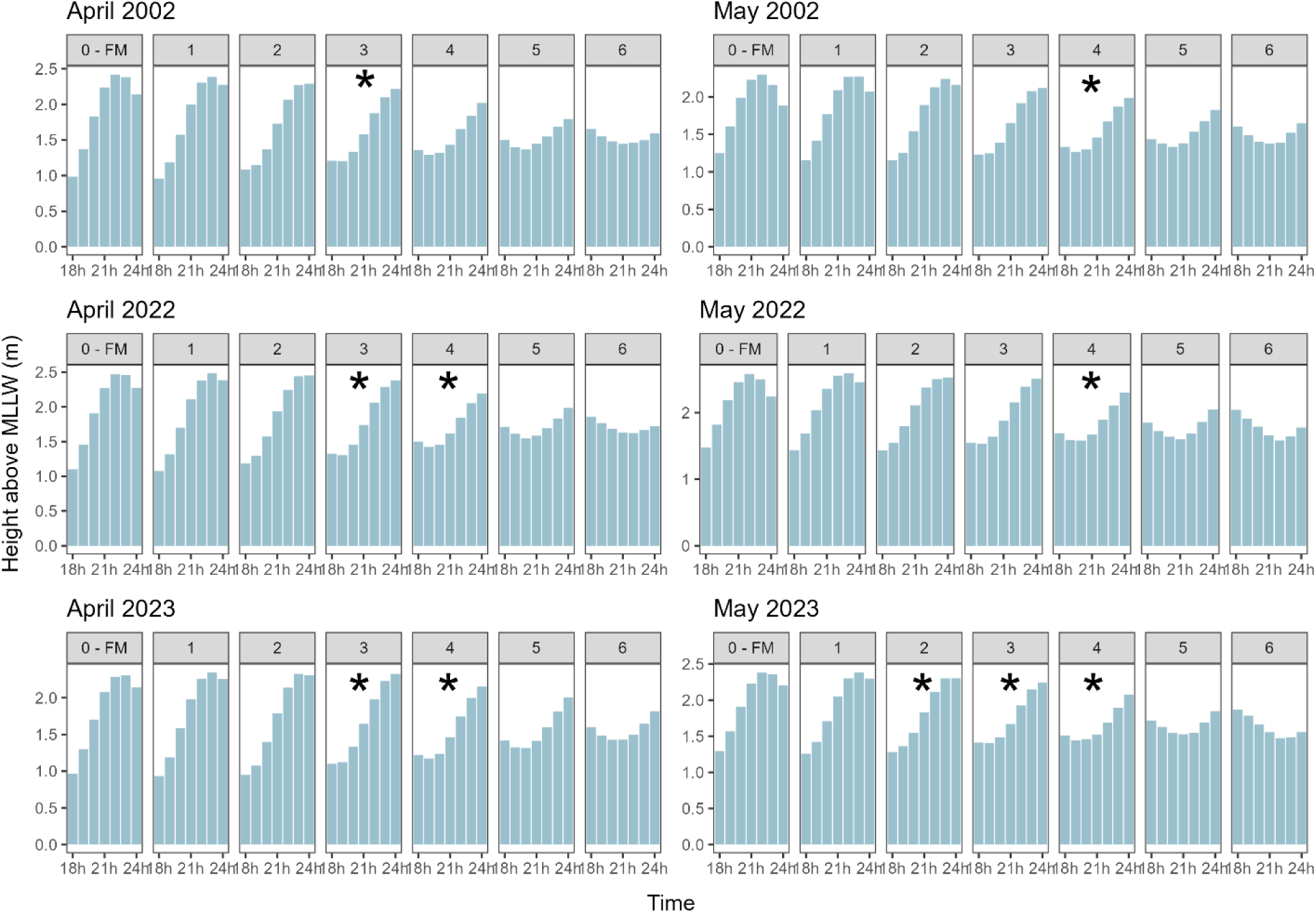
*Porites lobata* spawning observations in relation to tidal state. Hourly measurements of water level above mean low low water (MLLW) at Malakal Island, Palau (UHSLC Station #7) are shown from 18:00 to midnight on the night of the full moon and up to 6 nights afterward in each month. Asterix indicate nights when spawning was observed in 2002 (Penland et al. 2004), 2022, and 2023 (present study).

*Porites lobata* sperm had a small head with a pointed tip and wider base, plus a long flagellum (Figure 7). Sperm release resembled white smoke and occurred in pulses that were each several minutes apart (Figure 7). Female colonies released light pink-brown eggs that were solitary or in clusters of <5 eggs; the eggs were weakly buoyant and floated very slowly to the surface of the water (Figure 7). Mature oocytes could be observed under a dissecting microscope in tissue chips taken from gravid adult colonies prior to spawning (Figure 7). Eggs were spherical, homogeneous in terms of shape and size, and had a diameter of 233.7 ± 18.1 μm (mean ± stdev) (Figure 9). The algal symbiont concentration was 121 ± 26 cells per egg.

**Figure 7.**
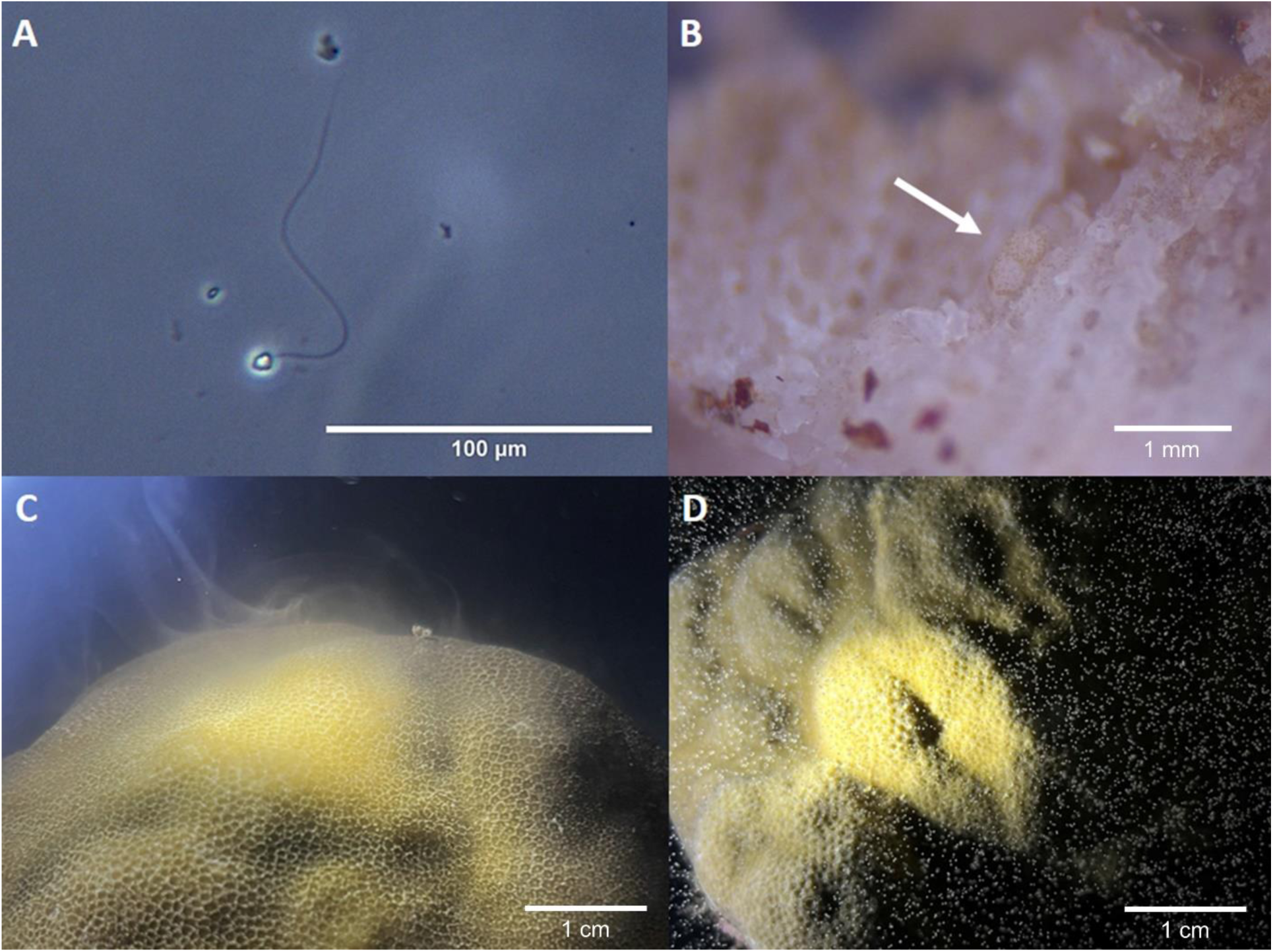
*Ex situ* spawning in *Porites lobata*. (A) Phase contrast light micrograph of sperm. (B) Mature oocytes observed in tissue sample from adult *P. lobata* colony. (C) Male beginning to release sperm with the appearance of white smoke. (D) Female releasing pink-brown eggs, some of which occur in small clusters.

### 3.3 Gamete age assays

Fertilization success was high in *Porites lobata* for the first 2 – 2.5 hours after spawning began for a given female (Figure 8). Eggs older than 2.5 hours had much lower fertilization success, with nearly zero fertilization 3.5 hours after spawning began. We also observed some variation in egg viability, with eggs from one female showing a precipitous decline in fertilization success earlier than other females tested (Figure 8). Sperm were observed actively swimming up to 4 hours after spawning began for a given male, indicating that sperm motility was likely not the limiting factor in fertilization success for eggs older than 2.5 hours.

**Figure 8.**
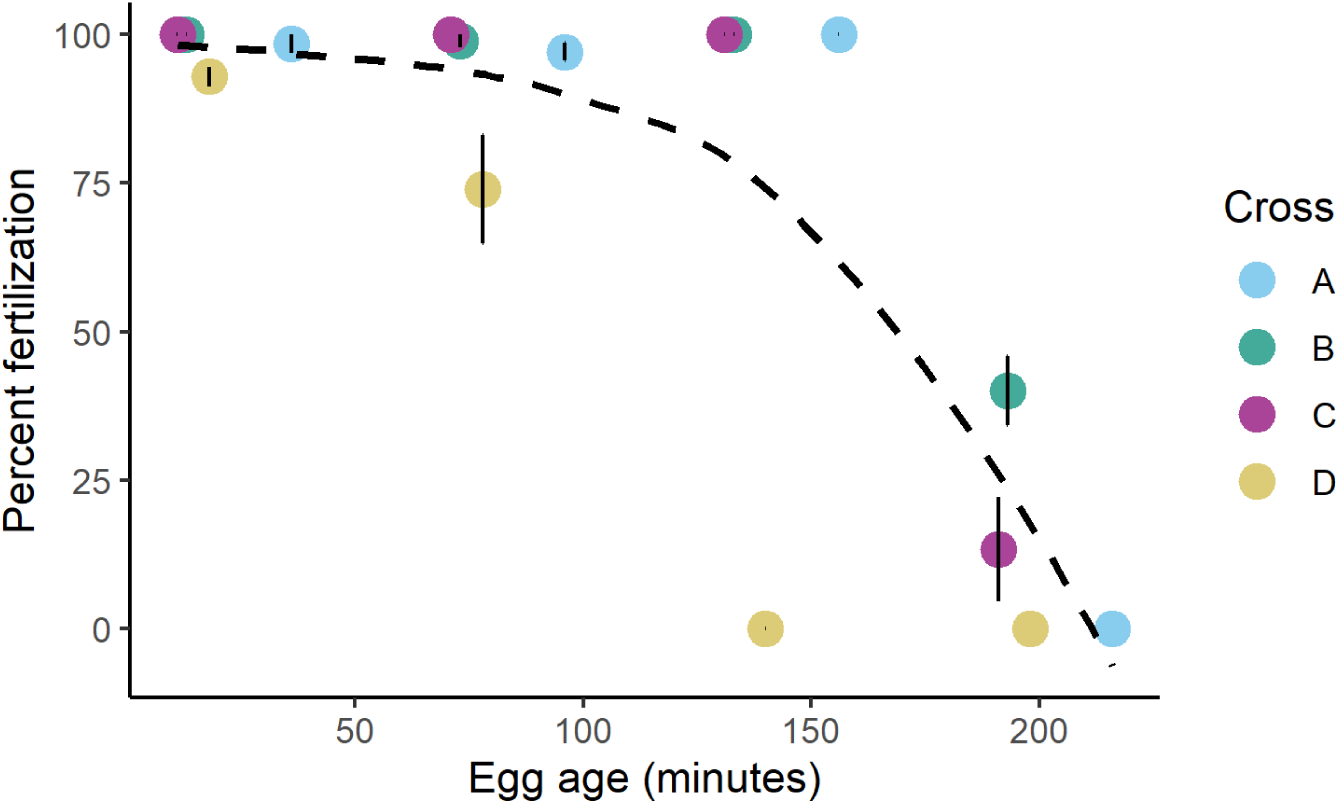
Increasing egg age reduces fertilization in *Porites lobata*. Each cross included eggs from one female and pooled sperm from 3 males. Error bars represent standard error. Dashed line represents best-fit curve. Colors represent the four different crosses of this assay. Each cross had three technical replicates per timepoint, where eggs from the same egg donor were added to three separate containers holding sperm from the three pooled sperm donors. Fertilization in each container was assessed separately and averaged to obtain the fertilization rate for a given cross and timepoint.

### 3.3 Embryological observations

The first cellular division (2-cell stage) was observed ∼1.5 hours after sperm and eggs were mixed. Cellular divisions rates varied among fertilized eggs, and batch crosses had individuals in multiple stages of embryonic development at the same time (Figure 9). Sub-samples of eggs maintained under a microscope in an air-conditioned room (27 °C) took longer to divide than eggs maintained at ambient temperature (∼31 °C), and many did not complete embryogenesis. Fertilization success and embryological development may sensitive to temperature in this species, though further dedicated study would be required to confirm our observation.

**Figure 9.**
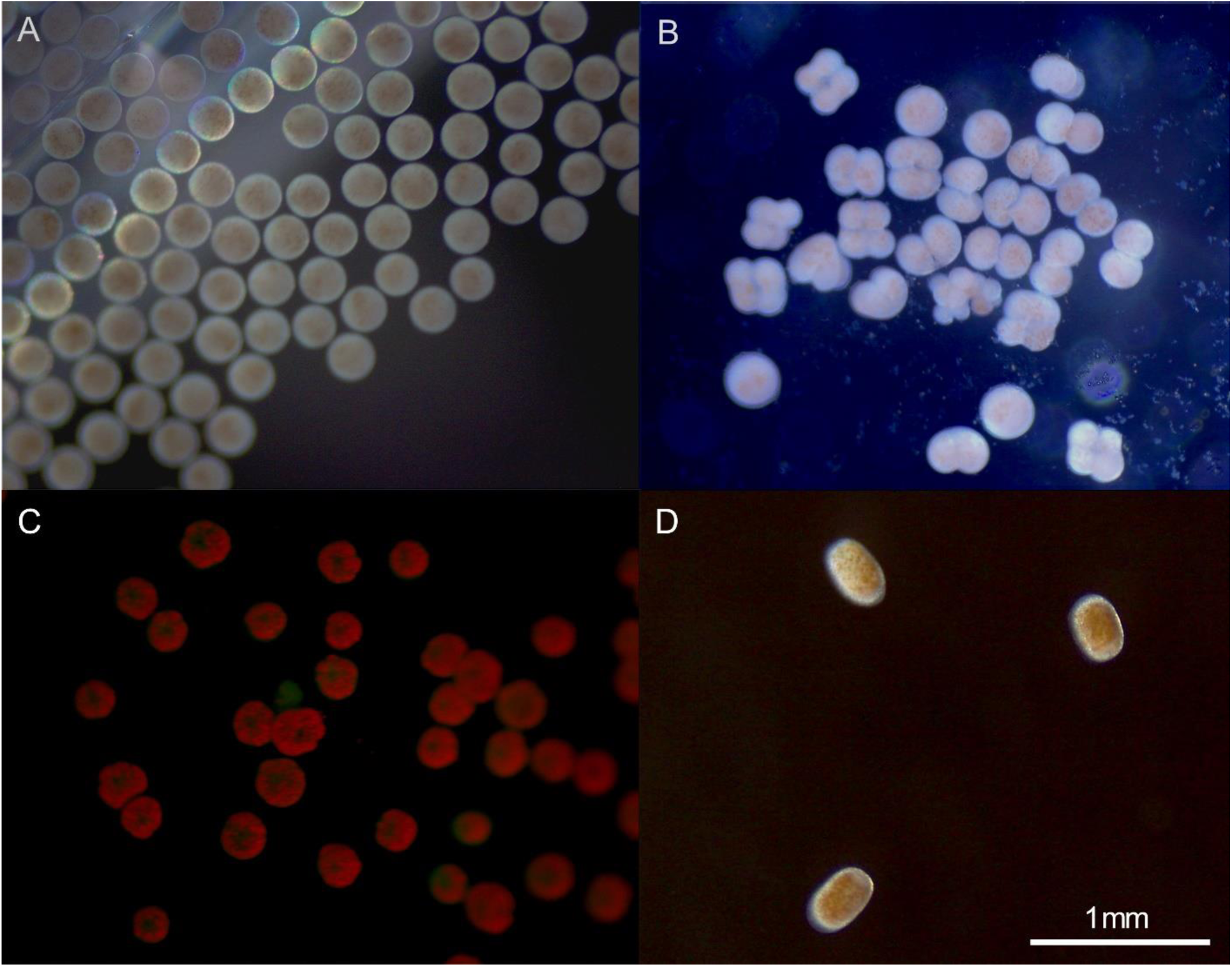
Development of *Porites lobata* larvae. (A) Representative picture of eggs directly after collection. (B) Fertilized embryos with dividing 2- and 4-cell stages. The first embryos observed to complete the first division did so roughly 90 minutes after gametes were mixed, and the same embryos completed their second division ∼40 minutes later when kept at ambient temperature. (C) Fluorescence in 3-day-old larvae. (D) 4-day-old larvae. Scale bar applies to all figure panels.

There were 207 ± 76 algae in 5-day-old *P. lobata* larvae. Larvae were elliptical, 312 ± 39 μm long and 254 ± 41 μm in the shorter diameter (Figure 9). Larvae were neutrally or negatively buoyant and were evenly distributed throughout the water column in culture containers 36 hours after fertilization. Most larvae had slow, directional or meandering swimming, although some barely swam at all in the still water cultures.

### 3.4 Settlement observations

Larvae older than 2 days began to display characteristic settlement behaviours, i.e., repeated exploration of the bottom of culture dishes or swimming in helices. The mean diameter of settled individuals was 580 ± 13 μm (n=19, range 486 – 675 μm) (Figure 10). Larvae readily settled on limestone tiles and glass microscope slides placed in plastic culture containers. Higher proportions of larvae settled on substrata where crushed CCA had been added (∼60 %), relative to substrata without CCA (<20 %). Some larvae in each culture settled in a narrow groove on the bottom of the culture containers, even without the addition of CCA as a settlement cue. Most larvae in settlement assays attached to the substratum within 24 hours, and calcification began two days after attachment (Figure 10). Tentacles were first observed 8 days after attachment to the substratum (Figure 10).

**Figure 10.**
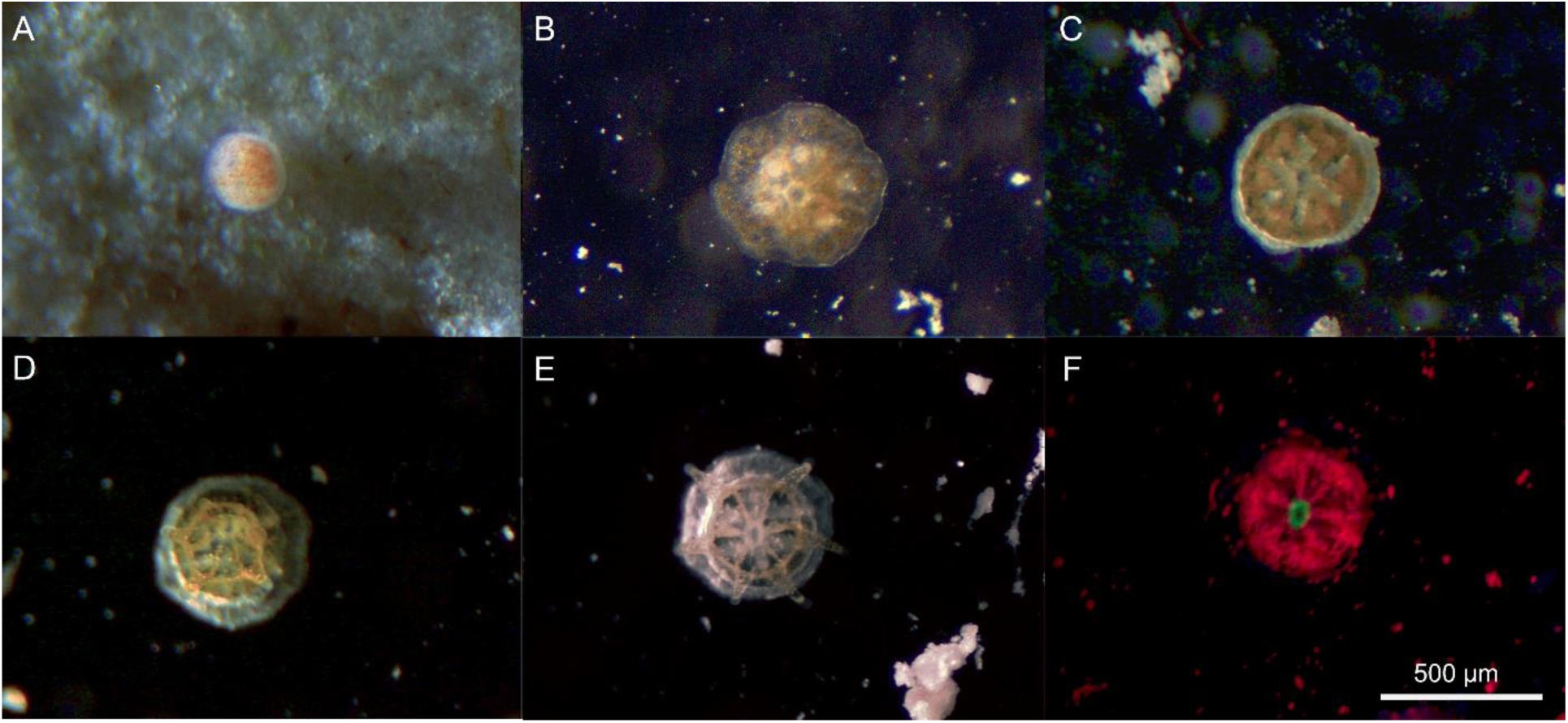
Settlement in *Porites lobata*. (A) Larva attached to limestone tile. (B) Settler 24 hours after attachment. (C) Settler 5 days after attachment. (D) Settler 7 days after attachment with rudimentary tentacles. (E) Settler 10 days after attachment. (F) Green fluorescence limited to polyp mouth in *P. lobata* settler 10 days after attachment. Scale bar applies to all figure panels.

## 4. Discussion

Our results confirmed that *Porites lobata* in Palau are gonochoric broadcast spawners with a spawning season spanning multiple months (Penland et al. 2004, Gouezo et al. 2020). Spawning occurred in the fourth and fifth lunar months in Palau, on nights with at least two hours of darkness between sunset and moonrise, on a rising tide. By relating spawning observations to environmental conditions, this study can aid in predicting *P. lobata* spawning events elsewhere in its broad geographic range.

We observed spawning in lunar months with a higher mean temperature than the previous lunar month. Rising water temperature is an important cue for coral gametogenesis and spawning across multiple spatial and temporal scales (Guest et al. 2005, Keith et al. 2016, Lin & Nozawa 2023). Increasing sea surface temperatures as a result of climate change could alter the timing of coral spawning (Paxton et al. 2016). However, we found spawning patterns in relation to environmental factors were remarkably consistent between our study (2022, 2023) and observations made 20 years prior (Penland et al. 2004). Water temperatures in shallow lagoons and coral reefs in Palau have risen ∼0.5° C in the last two decades (Colin 2018). This increase may not be large enough to cause a significant shift in *P. lobata* spawning timing, or the night of spawning within a given month might be determined by other factors, such as moonlight and tide. Corals rely on environmental cues on multiple temporal scales to “fine-tune” spawning timing and maximize fertilization success (Guest et al. 2008).

A period of darkness between sunset and moonrise appeared to be important for *P. lobata* spawning in this study. Exposure to light after sunset can disrupt spawning synchrony in corals, and in fact, darkness between sunset and moonrise is an important factor determining the night of spawning in multiple species (Kaniewska et al. 2015, Lin et al. 2021). Spawning mid-way between low and high tide is common among corals in Palau (Kenyon 1995). In bathymetrically complex areas such as our study sites in the Rock Island Southern Lagoon (Chelbacheb), spawning on an incoming tide could facilitate gamete mixing between reefs (Kenyon 1995).

Only a small proportion of coral colonies observed in this study spawned (2022: 17%, 2023: 35%) or had gametes in histological sections (2022: 18%). It is possible that we missed some spawning in April 2022, particularly for corals collected 4 dAFM. Nevertheless, colonies in this study were selected haphazardly (i.e., without regard to possession of gametes), so our observations in other months could be indicative of the proportion of individuals participating in spawning events *in situ*. A previous study in the eastern Pacific found only 30 – 67% of *P. lobata* colonies sampled had gonads (Glynn et al. 1994), though a study from Rapa Nui (Easter Island) reported very high (50-100 %) proportions of gravid colonies (Buck-Wiese et al. 2018). It is unclear why such a low proportion of colonies would be reproductive in a given year. No colonies were observed to switch sex between spawning years, so sex switching does not explain the low proportion of colonies spawning. Environmental stress, including bleaching, has carry-over effects on reproduction, leading to reduced fecundity, smaller oocytes, and a lower proportion of gravid colonies (Johnston et al. 2020, Leinbach et al. 2021). It is therefore possible that the low proportion of reproductive colonies in our study was a latent effect of environmental stress. The proportion of colonies spawning *ex situ* varied greatly among study sites, with no colonies from Merchechar spawning in either year. In contrast, 65% of colonies collected from Taoch released gametes in April or May 2023 (Supplementary Material). This could be related to differences in localized environmental conditions and recent history, or differences in the *P. lobata* populations found at each site.

One consistent observation in our data was that males commenced gamete release before females. Males spawning before females has also been reported for other gonochoric species (Hagman et al. 1998, Marhaver et al. 2015). Because we isolated colonies in individual containers to separate gametes, it is possible that females missed some chemical or environmental cues—such as the presence of sperm in the water column—that could have induced spawning sooner. A delay in gamete release *ex situ* has been reported when compared with concurrent field observations of the same species (Neely et al. 2021, O’Neil et al. 2021). Therefore, our observations should be confirmed by *in situ* spawning studies.

The wide time spans over which we observed spawning could serve as temporal pre-zygotic barriers in *P. lobata*. We observed spawning over two months per year, on multiple nights in each month, and over multiple hours on each night. While sperm swam actively for several hours, the time delay between sperm and egg release could result in sperm dilution and restricted fertilization rates *in situ* (Oliver & Babcock 1992). Furthermore, *P. lobata* eggs had high fertilization for only the first 2 – 2.5 hours after release. Other gonochoric coral species have similarly narrow windows for egg fertilization, which may serve as a reproductive barrier for sympatric species (Levitan et al. 2004). It is well-documented that *P. lobata* has high genotypic diversity and multiple cryptic lineages (Boulay et al. 2013, Forsman et al. 2015, Schweinsberg et al. 2016, Rivera et al. 2022). Future studies should examine the potential for spatial and temporal reproductive isolation among lineages.

Previous studies have assumed a long pelagic larval duration for *P. lobata* because of its vertical symbiont transmission (Field 1988, Baums et al. 2012). However, we observed competency to settle in *P. lobata* larvae just two days post fertilization. While individual larvae have been kept in culture for up to 53 days in a prior study (Field 1988), many individuals in our study settled on culture dishes in the absence of a settlement cue, suggesting “desperate” settlement. Factors leading to “desperate” behaviour in symbiotic larvae could be related to algal physiology and should be further explored. Our observations indicate that the vast majority of individuals settle shortly after achieving competency. Besides the abbreviated larval period, neutral or negative buoyancy was observed in most larvae within 24 hours of fertilization, with many individuals distributed throughout the water column or swimming at the bottom of the culture dishes. Variations in dispersal depth could lead to differences in dispersal distance for *P. lobata* larvae, since currents are slower, less directional, and more turbulent in the benthic boundary layer (Vogel 1996).

Palau’s Rock Island Southern Lagoon (Chelbacheb) is characterized by restricted circulation and long water residence times (Golbuu et al. 2016). Most reefs in this area are likely to be self-seeding (Golbuu et al. 2012, 2016). With a pelagic larval duration of just 2 days, *P. lobata* would be expected to have highly localized recruitment (Golbuu et al. 2012). Nevertheless, model results indicate that coral dispersal and recruitment are heavily influenced by variations in current direction and wind speed (Gouezo et al. 2021). Molecular studies of population genetic structure have suggested limited dispersal for *P. lobata* on multiple scales— between semi-isolated lagoons and outer patch and barrier reefs in Palau (Rivera et al. 2022), between the Hawaiʻian Islands and Johnston Atoll (Polato et al. 2010), and at the largest scale, across the East Pacific Barrier (Baums et al. 2012). Future studies should address how variations in pelagic larval duration and oceanographic conditions influence dispersal of *P. lobata*.

Overall, the results of this study contribute to the body of knowledge on the spawning, embryonic development, and larval settlement for a key reef-building coral species. We have been able to demonstrate high fertilization success and high settlement rates can be achieved with methods commonly practiced in *ex situ* reproductive studies. Furthermore, the short pelagic larval duration and high settlement achieved in our study, together with the high thermal tolerance and genetic diversity of *P. lobata* shown elsewhere, make it a promising candidate for *ex situ* assisted reproduction programs (Guest et al. 2023). By reporting specific timing for spawning events, and determining the optimal range for introducing gametes in *in vitro* fertilization procedures, our study should further enable restoration practitioners to include this species in future initiatives and optimize their fertilization success. Finally, development of a coral spawning calendar for Palau would rapidly accelerate research and enhance the capacity for larval propagation in this archipelago with additional coral species (e.g., Baird et al. 2022).

## Acknowledgments

We are grateful to Joy Shmull-Sam and the Palau International Coral Reef Center staff for facilitating field and lab work for this study in Palau. This material is based upon work supported by the National Science Foundation under Grant Nos. OCE-2048678 (KMK) and OCE-2048589 (SWD). Sample collection was authorized by scientific research permits issued by the Palau Ministry of Natural Resources, Environment, and Tourism (RE-22-17, RE-22-24, RE-23-09) and Koror State Government (permit nos. 72, 78, and 83) to KMK.

## Author contributions

Study design – MJB, KMK, CGBG, JD, SWD; Data collection and spawning observations – MJB, MA, KMK, CGBG; Histology – JD, AR, DH; Data analysis – MJB, CGBG, JD, AR, DH; Manuscript drafting – MJB, KMK, CGBG, JD; Manuscript editing – all.

