## Supplementary Material for "*Ex situ* spawning, larval development, and settlement in the massive reef-building coral *Porites lobata* in Palau"

**Supplementary Table 1.** Summary of the number of colonies collected and observed in *ex situ* aquariums during the spawning windows in April and May 2023, and the proportion of colonies observed releasing gametes in either month by site.

| Site | Latitude (N) | Longitude (E) | No. colonies collected | No. colonies spawning | % of colonies spawning |
|---|---|---|---|---|---|
| Merchechar | 7.16162 | 134.34923 | 9 | 0 | 0% |
| Ngelsibel | 7.18686 | 134.34697 | 10 | 2 | 20% |
| Ngerur | 7.3672 | 134.4298 | 11 | 1 | 9% |
| Outer Taoch | 7.27193 | 134.38115 | 29 | 11 | 38% |
| Risong | 7.30971 | 134.47661 | 13 | 3 | 23% |
| Taoch | 7.26723 | 134.38722 | 26 | 17 | 65% |

### Preliminary sperm cryopreservation trials

Given the potential for reproductive isolation between populations of *P. lobata* due to their broad distribution, cryopreservation of sperm may become an important technique to support sexual propagation and increase genetic diversity in this species (Hagedorn et al., 2017; Baums et al., 2012).

Preliminary cryopreservation tests were conducted using sperm from a subset of males (n=10 males). Sperm were collected directly from the surface of coral colonies during release, divided into 5-10 aliquots (1 mL each), and frozen in liquid nitrogen using the design by Zuchowicz et al. (2021). Sperm samples were frozen without (n=10/10 males) or with (n=5/10 males) the cryoprotectant dimethyl sulfoxide (DMSO, 10% solution) to test how it affected sperm integrity and competency. We also concentrated larger samples of sperm (45–100 mL) by centrifugation at 3000 x *g* and 7000 x *g* for 5 minutes and resuspending the pellet in 1 mL FSW prior to cryopreservation (n=3 males). Sperm aliquots were kept frozen for 24 hours up to ~1 month. After thawing, sperm motility was assessed in a subset of samples (n=5) using a phase contrast microscope at 10–40x magnification. Sperm competency was tested by mixing

1–3 mL of thawed sperm from 1–2 males with viable, unfertilized, eggs from 1–2 females in triplicate (n=10 crosses with sperm samples containing DMSO, 11 crosses without DMSO).

Intact sperm cells were observed in all thawed samples. Generally, the addition of DMSO resulted in visibly higher (~10x) recovery of intact sperm cells, but exact concentrations were not quantified. Centrifugation at 3000 x *g* and 7000 x *g* visibly increased sperm concentrations in 1 mL aliquots, but centrifugation at 7000 x *g* resulted in tail breakage and reduced sperm motility. Successful fertilization (>50% eggs fertilised) was observed in two crosses using cryopreserved sperm without the addition of DMSO (all replicates). Fertilization was also observed in one cross with DMSO (all replicates). DMSO did not appear to reduce fertilization rates; however, eggs fertilised by sperm stored with DMSO had irregular development and died after 2 – 4 hours, suggesting that 10% DMSO may be toxic to *P. lobata* eggs or hinder larval development.

Our preliminary fertilization trials with cryopreserved sperm resulted in limited fertilization success (1/11 crosses with DMSO, 2/10 crosses without DMSO), suggesting that further optimisation of this technique is necessary for *P. lobata*. While the use of DMSO as a cryoprotectant increased the yields of intact sperm after thawing, it reduced the survival of fertilised eggs. For future work, we recommend ultrastructure examination to understand the underlying mechanisms of unsuccessful fertilization. DMSO should be added while freezing sperm, but sperm should be cleaned of DMSO prior to egg fertilization by using multiple cycles of centrifugation (3,000 x *g*) and rinsing the pellet with FSW.
